# Molecular Basis of Behavioral Diversity in a Sibling Species Trio

**DOI:** 10.64898/2026.02.23.707296

**Authors:** Cynthia M. Chai

**Affiliations:** Department of Biological Sciences, Columbia University, 1212 Amsterdam Ave, New York, NY 10027, USA

**Keywords:** Brain evolution, Gene expression, Sibling species, Speciation, Bioinformatic theory

## Abstract

The brain is the main controller of animal behaviors. During the speciation process, divergent behaviors can arise between new sibling species. However, the comprehensive molecular changes in the brain that give rise to these emergent behavioral differences are not well understood. Here, I present a comparative analysis of gene expression differences across the entire nervous system at region-specific resolution between a trio of closely-related *Drosophila* sibling species representing two consecutive speciation events. My analysis revealed strongest patterns of molecular conservation across speciation events in the brain region that executes movement relative to regions associated with other neurobehavioral functions. The RNA-seq dataset and analytical strategy discussed here also establishes a new resource to accelerate mechanistic comparative neuroscience studies in this genetically accessible yet understudied sibling species trio. As genomic resources expand (BLAXTER *et al*. 2025), iteratively exploring these patterns across ever larger swaths of animal phylogeny is poised to reveal general principles of brain functional molecular evolution.

## INTRODUCTION

Darwin’s observation of the myriad forms of finch beaks in the Galapagos Islands provides a classic example of the rich biodiversity that arises as animals adapt to novel ecological niches (DARWIN 1909). Genomic variation impacting essential behaviors, such as foraging and mating, can drive reproductive isolation between populations that exhibit differential responses, ultimately resulting in speciation. However, the proximate mechanisms in the nervous system underlying behavioral divergence between new sibling species are only sparsely documented. Recent studies using closely-related sibling species have reported either cell-type specific transcriptomic differences between homologous neurons or transcriptomic changes in a subset of brain regions between species or whole-brain bulk changes between species pairs (MONTGOMERY *et al*. 2021; KAUTT *et al*. 2025; LEE *et al*. 2025; WALSH *et al*. 2025). Here I take a complementary approach by providing a global view of comparative gene expression changes across the entire nervous system at region-specific resolution in a trio of closely-related sibling species representing two consecutive speciation events.

## RESULTS AND DISCUSSION

Like *Drosophila melanogaster, Drosophila simulans*, which diverged from *D. melanogaster* around 3 million years ago, is a cosmopolitan species that has colonized diverse habitats (DEAN AND BALLARD 2004; DAVID *et al*. 2007; GARRIGAN *et al*. 2012) (Figure 1A). In contrast, *Drosophila mauritiana* is endemic to the volcanic islands of Mauritius and diverged from *D. simulans* around 250,000 years ago (DAVID *et al*. 2007; GARRIGAN *et al*. 2012) (Figure 1A). In the *melanogaster* species subgroup, a premier model clade for studying neurobehavioral evolution (JOURJINE AND HOEKSTRA 2021), the island endemic *D. mauritiana* has not been studied in regard to the neural basis of behavioral evolution. The *Drosophila* nervous system can be partitioned into three anatomical regions associated with generally distinct hierarchical neurobehavioral functions: visual sensory (optic lobes), higher-order processing (central brain), and movement (ventral nerve cord) (Figure 1B). By comparing region-specific bulk transcriptomic profiles between all three sibling species (Figure 1C and S1A,B), I first sought to determine the relative levels of molecular divergence underlying distinct neurobehavioral functions as indicated by the extent of inter-specific differential gene expression.

**Figure 1.**
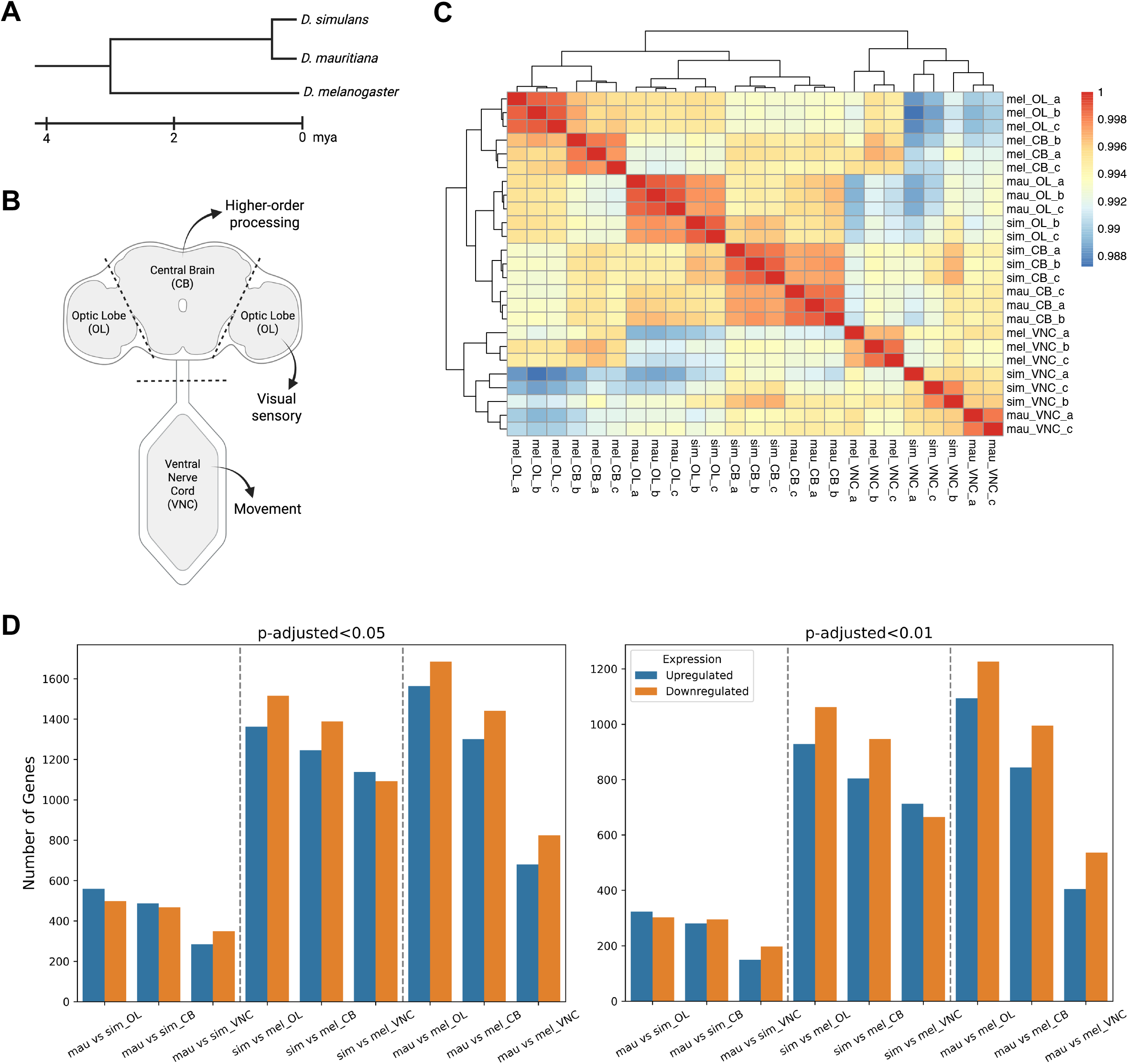
Comparative brain transcriptomics in a sibling species trio representing two consecutive speciation events reveals brain regional patterns of molecular evolution. (A)Phylogeny of *Drosophila melanogaster, Drosophila simulans*, and *Drosophila mauritiana* sibling species (GARRIGAN *et al*. 2012). (B)Schematic of dissected *Drosophila* brain regions labeled with their associated general neurobehavioral functions (ASO *et al*. 2014; NÉRIEC AND DESPLAN 2016; COURT *et al*. 2020; HULSE *et al*. 2021). (C)Hierarchical clustering heatmap of correlation matrix for bulk RNA-seq samples (25 samples). (D)Number of significantly differentially expressed genes, p-adjusted<0.05 (Left) and 0.01 (Right), for each species pairwise test per brain region. Number of genes with non-zero total read count: optic lobes (13,012), central brain (13,325), and ventral nerve cord (13,246).

As RNA-seq reads were mapped to the *D. melanogaster* reference genome for all sibling species (Figure S2), *D. simulans* and *D. mauritiana* species-specific genes are therefore excluded from this study’s analyses (MYERS *et al*. 2000). Across consecutive speciation events, the ventral nerve cord, which primarily orchestrates movement, exhibited the lowest number of significantly differentially expressed genes between sibling species (Figure 1C and 1D). Conversely, the optic lobes, a major sensory modality in *Drosophila*, was consistently most divergent in gene expression across speciation events (Figure 1D). In other words, how a fly responds to an external challenge, rather than its ability to respond, appears more likely to differ between speciated populations at the molecular level. Interestingly, comparative single-cell transcriptomic analysis of three *Caenorhabditis* nematode species spanning an evolutionary divergence time of 45 million years identified strongest conservation of orthologous gene expression in motor neuron classes relative to other neuron functional categories (TOKER *et al*. 2025).

As the comparative RNA-seq experimental design here comprises a trio of sibling species mapped to the same reference genome rather than just a species pair, phylogeny can thus be used to constrain the number of candidate neuronal genes that could underlie inter-specific behavioral differences. To concretely illustrate this, I introduce a locomotor activity behavior difference between sibling species (Figure 2A,B). As *D. simulans* and *D. mauritiana*, which are members of the *simulans* clade, consistently exhibited the same decreased locomotor activity phenotype relative to *D. melanogaster*, neuronal genes that are differentially expressed in both *D. simulans* (Set A) and *D. mauritiana* (Set B) compared to *D. melanogaster* are more likely to contribute to the behavioral phenotype (Figure 2C and 2D). The intersection of both sets (A ∩ B) results in a 41-52% reduction in candidate gene set size relative to A and B across brain regions (Figure 2D). Neuronal genes that are significantly differentially expressed between *D. mauritiana* and *D. simulans* are not likely to contribute to the shared decreased locomotor activity phenotype. Thus, subsequently obtaining the set difference of A ∩ B and the set of differentially expressed genes between *D. mauritiana* and *D. simulans* (Set C) results in a marginal further decrease in candidate neuronal gene set size ((A ∩ B) \ C) (Figure 2C and 2D).

**Figure 2.**
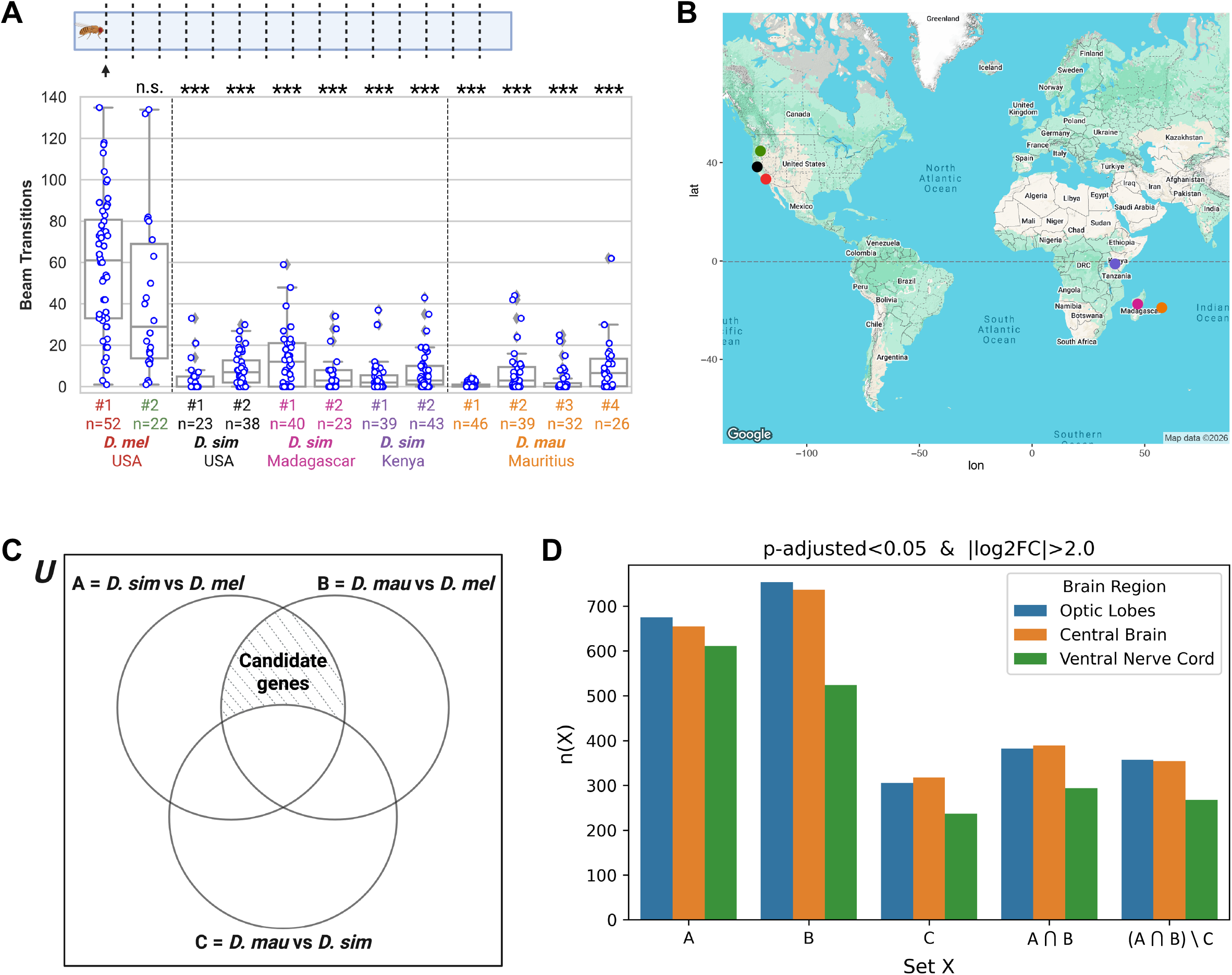
Parallel analysis of species trios at both the genotype and phenotype levels significantly reduces the search space for the causal genomic variation underlying behavioral evolution. (A) (Top) Schematic of locomotor activity assay with black arrow indicating an infrared beam intersecting the glass assay tube. (Bottom) Locomotor activity behavior for wild-type *Drosophila* sibling species lines. The total number of infrared beam transitions during the 30-min assay duration is a proxy for locomotor activity levels. n=number of flies. Kruskal-Wallis followed by Dunn’s post hoc with Holm correction, n.s.=no significance, ***p<0.001. (B)World map showing geographical origins of *Drosophila* species lines assayed in Figure 2A. Dot color corresponds to species x-axis label color in Figure 2A. (C)Set operations strategy to constrain the number of candidate neuronal genes underlying observed inter-specific differences in locomotor activity behavior. The universal set (*U*) is defined as the finite set of all annotated genes in the *D. melanogaster* reference genome. (D)Comparison of brain region-specific candidate neuronal gene set sizes before and after executing set operations strategy in (C). n(X) denotes the cardinality of a given set X (i.e. the number of differentially expressed genes in set X).

The brain region-specific transcriptomics dataset and relevant set operations strategies presented here generalizes and can be applied to all behavioral phenotype combinations examined in this sibling species trio (Figure 3A and Figure S3A,B,C,D). Moreover, inclusion of two consecutive speciation events in the experimental design facilitates the inference of corresponding molecular evolution scenarios underlying behavioral divergence (Figure 3B). In the absence of *a priori* knowledge regarding the identity of and changes in neural circuitry underlying inter-specific behavioral differences, spatial analysis of candidate differentially expressed genetic elements reported in this study can provide a path towards uncovering the neuronal sites of evolutionary innovation. Tracing the specific tunable steps leading to the emergence of new species could enable us to better understand and predict Life’s trajectory in an ever-changing biosphere.

**Figure 3.**
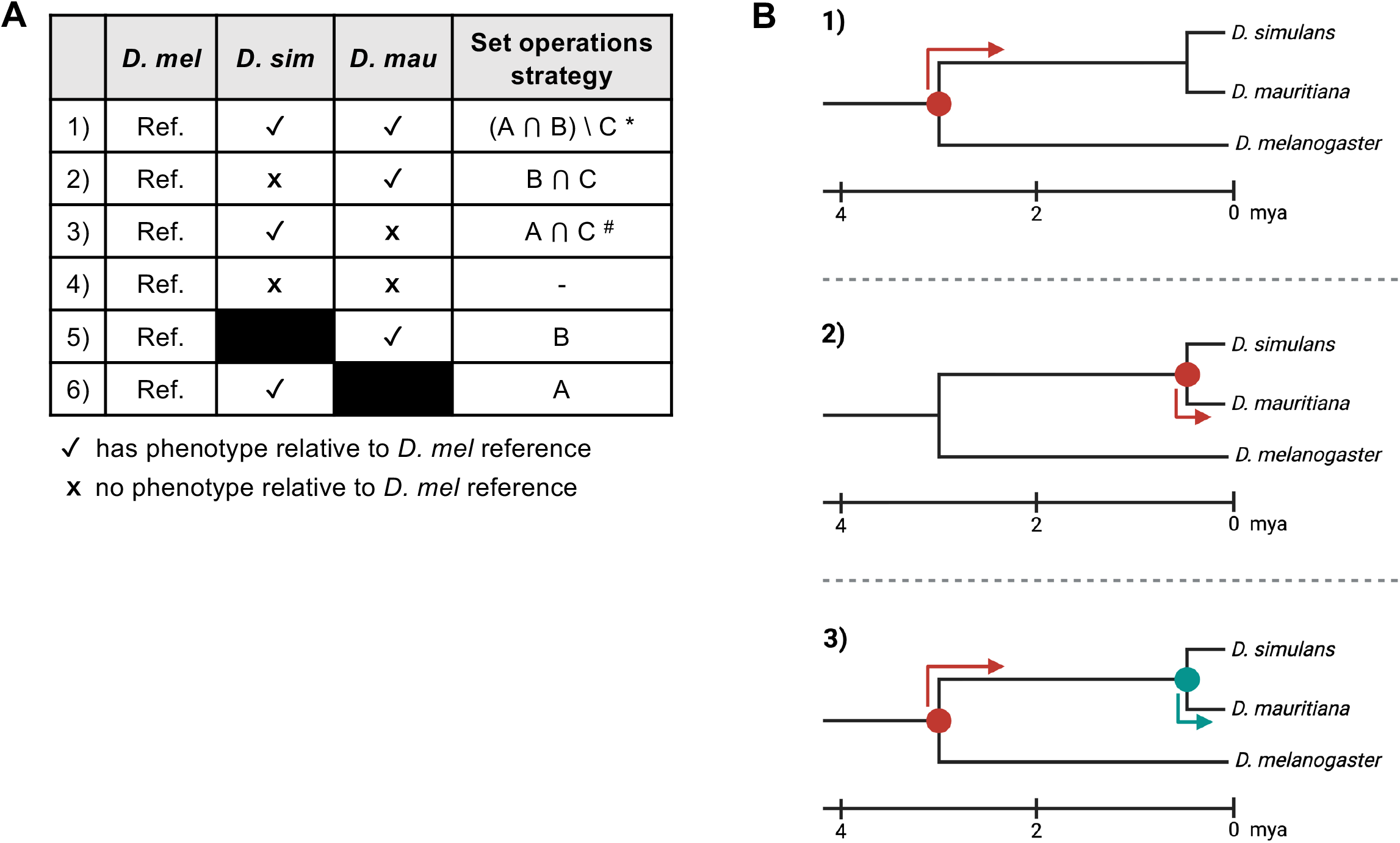
Mapping behavioral phenotype combinations to species phylogeny provides a generalizable approach to constrain the number of candidate neuronal genes underlying behavioral differences between this sibling species trio. (A)General behavioral phenotype combinations and corresponding set operations strategies to constrain candidate neuronal gene set sizes. See SI Software for code implementation. See Figure 3B legend for notes related to special cases denoted by * and ^#^. (B)Molecular evolution scenarios corresponding to sibling species phenotypic outcomes in Figure 3A. For each scenario, the circle indicates the relevant speciation event and the arrow indicates the branch along which the causal genomic variation could have occurred. For 3), green indicates an independent change subsequent to the initial change in red that reverts the *D. mauritiana* behavior to the *D. melanogaster* behavior. Figure 3A special cases: (C)Working assumption: The valence of both the *D. simulans* and *D. mauritiana* behavioral phenotypes relative to the *D. melanogaster* reference is in the same direction. If not, then the set operations strategy is A or B. ^#^ Working assumption: The subsequent genomic change impacts the same gene(s) impacted by the initial change. If not, then the set operations strategy is A or C.

## Supporting information

Software S1

Software S2

Software S3

Software S4

Software S5

## DATA AND SOFTWARE AVAILABILITY

Datasets and software code needed to re-analyze the results of this study are available in the manuscript supplement. All raw sequencing read files will be deposited in a publicly accessible online repository upon publication.

## AUTHOR CONTRIBUTIONS

C.M.C. conceptualized and designed the study, performed all experiments, analyzed RNA-seq data, acquired resources and funding, prepared the figures and wrote the paper. Generative AI and AI-assisted technologies were not utilized during the preparation of this manuscript.

## ACKNOWLEDGMENTS

The Janelia Quantitative Genomics Team performed RNA-seq sample preparation and sequencing. P. Andolfatto provided fly food, some fly species stocks, and equipment.

## FIGURE LEGENDS

## MATERIALS AND METHODS

### Animal stocks and maintenance

*Drosophila* fruit flies were raised on molasses-based fly food Recipe J in narrow vials (Lab Express, MI). The following *Drosophila* lines were used in this study:

| Species | Stock ID | Origin | Source | Figure 2A |
| --- | --- | --- | --- | --- |
| *Drosophila melanogaster* | Dickinson Lab (DL) | Pasadena, California, USA | Collected by M.H. Dickinson | D. mel USA #1 *** |
| *Drosophila melanogaster* | Oregon-R | Oregon, USA | Bloomington *Drosophila* Stock Center, Indiana University Bloomington | D. mel USA #2 |
| *Drosophila simulans* | D. sim 14021-0251.194 | Winters, California, USA | National *Drosophila* Species Stock Center, Cornell University | D. sim USA #1 *** # |
| *Drosophila simulans* | D. sim 14021-0251.216 | Winters, California, USA | National *Drosophila* Species Stock Center, Cornell University | D. sim USA #2 |
| *Drosophila simulans* | MD06 | Madagascar | Collected by B. Ballard in 2002 | D. sim Madagascar #1 |
| *Drosophila simulans* | MD221 | Madagascar | Collected by B. Ballard in 2002 | D. sim Madagascar #2 |
| *Drosophila simulans* | NS37 | Nairobi, Kenya | Collected by P. Andolfatto in 2006 | D. sim Kenya #1 |
| *Drosophila simulans* | NS67 | Nairobi, Kenya | Collected by P. Andolfatto in 2006 | D. sim Kenya #2 |
| *Drosophila mauritiana* | D. mau(1) SD 14021-0241.01 | Mauritius | National *Drosophila* Species Stock Center, Cornell University | D. mau Mauritius #1 *** |
| *Drosophila mauritiana* | R10 | Mauritius | - | D. mau Mauritius #2 |
| *Drosophila mauritiana* | R13 | Mauritius | - | D. mau Mauritius #3 |
| *Drosophila mauritiana* | R27 | Mauritius | - | D. mau Mauritius #4 |

\*\*\* Lines used to generate bulk RNA-seq dataset.

^#^ I incidentally discovered that the *D. simulans* 14021-0251.194 line (*D. simulans* USA Line #1) harbors a 4.6 kb retrotransposon insertion within the first exon of *Irk3* (inwardly rectifying potassium channel), likely resulting in a null allele. Targeted PCR-based genotyping of the *Irk3* locus indicates that the insertion is a Long Terminal Repeat (LTR) retrotransposon belonging to the Ty1-*copia*/Pseudoviridae family (classification by InterPro (BLUM *et al*. 2024), EMBL-EBI). Correspondingly, no *Irk3* transcript expression was detected for *D. simulans* in the RNA-seq dataset. Targeted PCR-based genotyping of the *Irk3* locus confirmed that this structural variant was not present in the second *D. simulans* 14021-0251.216 line isolated from Winters, California that was also assayed for behavior in this study. Of relevance, the retrotransposon-mediated *Irk3* knockout in *D. simulans* USA Line #1 does not result in a locomotor activity behavior difference relative to other *D. simulans* lines (Figure 2A).

### Locomotor activity behavior assay

Three to five-day old adult female flies raised at 22-23° C and 30-35% humidity on a 12-hr light/12-hr dark cycle were assayed. Locomotor activity behavior assays were performed using a DAM5H *Drosophila* activity monitor device (TriKinetics Inc., Waltham MA) connected to a computer running the DAMSystem3 data collection program. Each monitor device can assay up to 32 flies in parallel. Fly movement within each glass assay tube was recorded as the interruption of fifteen equally spaced infrared beams. Each assay tube was sealed on each end using small plugs fashioned from Kimwipes. The number of times each fly moved from one beam to another (i.e. beam transitions) was tallied across the entire trial duration for each animal.

Flies were cold anesthetized on ice prior to loading into glass assay tubes and were given 15 minutes of recovery time in the device followed by 30 minutes of locomotor activity recording time. All assays were performed during the four-hour duration before incubator lights out. Data collection was performed at 24-25°C and 35% humidity. Representatives of each *Drosophila* sibling species were always assayed on each day of experiments. Data across all trials were pooled for statistical analysis. Statistical analyses were performed using the *SciPy* and *scikit-posthocs* packages in Python.

### Cross-species brain transcriptomics

Brains were dissected from three to five-day old adult female flies raised at 22°C and 50% humidity on a 16-hr light/8-hr dark cycle. For each sibling species, ten whole brains were dissected per sample (a, b, or c) and their optic lobes, central brain, and ventral nerve cords were sub-dissected and pooled in Trizol (Invitrogen #15596020) for lysis followed by total RNA extraction. cDNA was generated using the same buffer and enzymes as described previously (CEMBROWSKI *et al*. 2018).

For each sample, 1 ng of purified total RNA was reverse transcribed using:

- RT primer (5’-TCG TCG GCA GCG TCA GAT GTG TAT AAG AGA CAG TTT TTT TTT TTT TTT TTT TTT TTT TTT TTT VN) and
- TSO primer (/5’Biosg/GTCTCGTGGGCTCGGAGATGTGTATAAGAGACArGrGrG)

PCR amplification was performed using primers:

- R1: 5’-TCG TCG GCA GCG TCA GAT GTG TAT AAG AGA CAG
- R2: 5’-GTC TCG TGG GCT CGG AGA TGT GTA TAA GAG ACA G

cDNA was purified with Ampure XP Beads (0.6x ratio; Beckman Coulter) followed by quantification using Qubit and bioanalyzer. Sequencing libraries were generated using a Nextera XT kit (Illumina #15027987) and sequenced on an Illumina Nextseq 2000 platform, with read lengths of 75 bp for read 1 and read 2; 10 bp for i7 and i5. Sequencing adapters were trimmed from reads using *Cutadapt* prior to alignment to the well-annotated *D. melanogaster* reference genome with STAR (DOBIN *et al*. 2013). The resulting transcript alignments were then passed to RSEM to generate gene expression counts (LI AND DEWEY 2011). Differential gene expression analysis was performed using DESeq2 in R. Two pre-identified outlier samples, ‘mau_VNC_b’ and ‘sim_OL_a’, are removed for all differential gene expression analyses (Figure S1).

### Targeted PCR-based genotyping

Flies were homogenized with a pipette tip and genomic DNA was extracted using a QIAwave DNA Blood & Tissue Kit (Qiagen #69554). The *Irk3* locus was PCR amplified using genomic DNA as template, and the PCR product was purified using a DNA Clean & Concentrator-5 Kit (Zymo #D4003T). The following genotyping primers (Integrated DNA Technologies, 25 nmole DNA oligonucleotides) were used: ACATATACGAGCGTGACATGAC (for) and GATAGCTTCATGCGCGAGTTC (rev). Amplicon sequencing was performed by Plasmidsaurus, Inc.

## SUPPLEMENTAL INFORMATION

### SUPPLEMENTAL FIGURE LEGENDS

**Figure S1.**
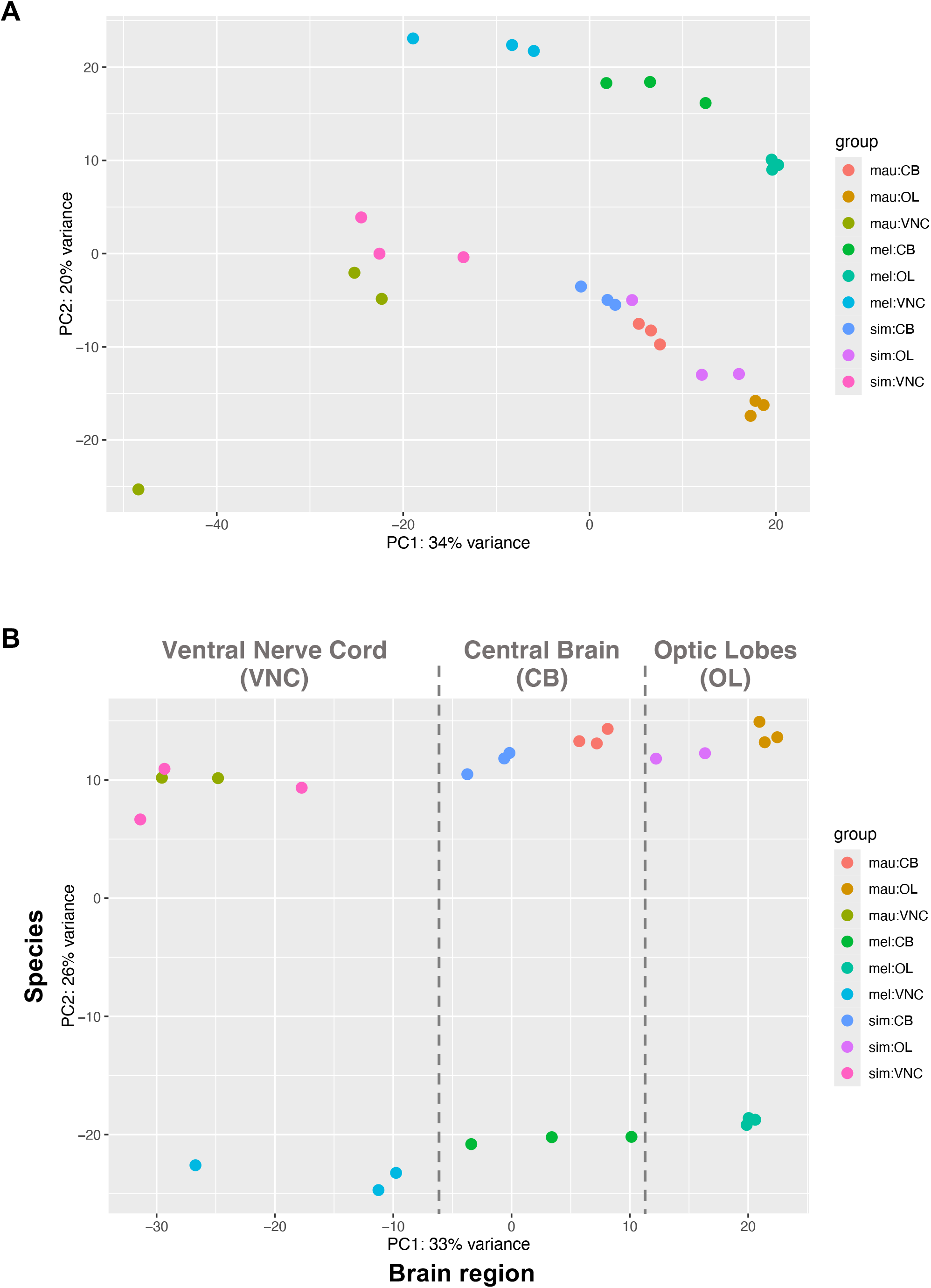
Brain region-specific bulk RNA-seq for three closely related *Drosophila* sibling species. Related to Figure 1. (A)Principal component analysis of all bulk RNA-seq samples (27 samples). (B)Principal component analysis of bulk RNA-seq samples (25 samples). Two pre-identified outliers, one *D. mauritiana* ventral nerve cord sample and one *D. simulans* optic lobe sample, were removed here and for all subsequent analyses.

**Figure S2.**
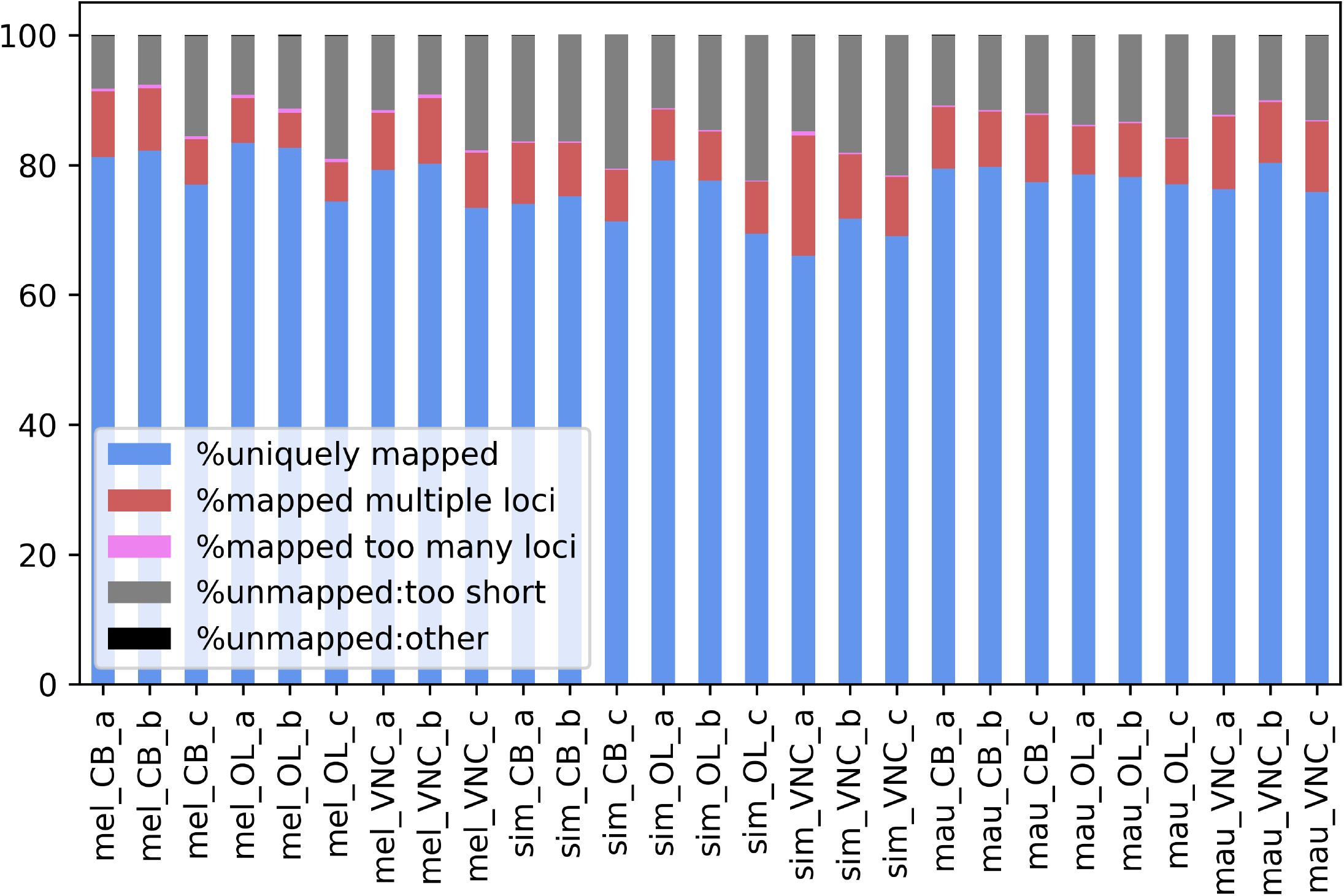
STAR RNA-seq aligner performance for read mapping of all RNA-seq samples to the *D. melanogaster* reference genome (Dobin *et al*., *Bioinformatics* 2013). Related to Figure 1. Percent ranges of uniquely mapped reads per species: *D. melanogaster* (73.5-83.4%), *D. simulans* (66.1-80.7%), and *D. mauritiana* (75.9-80.3%).

**Figure S3.**
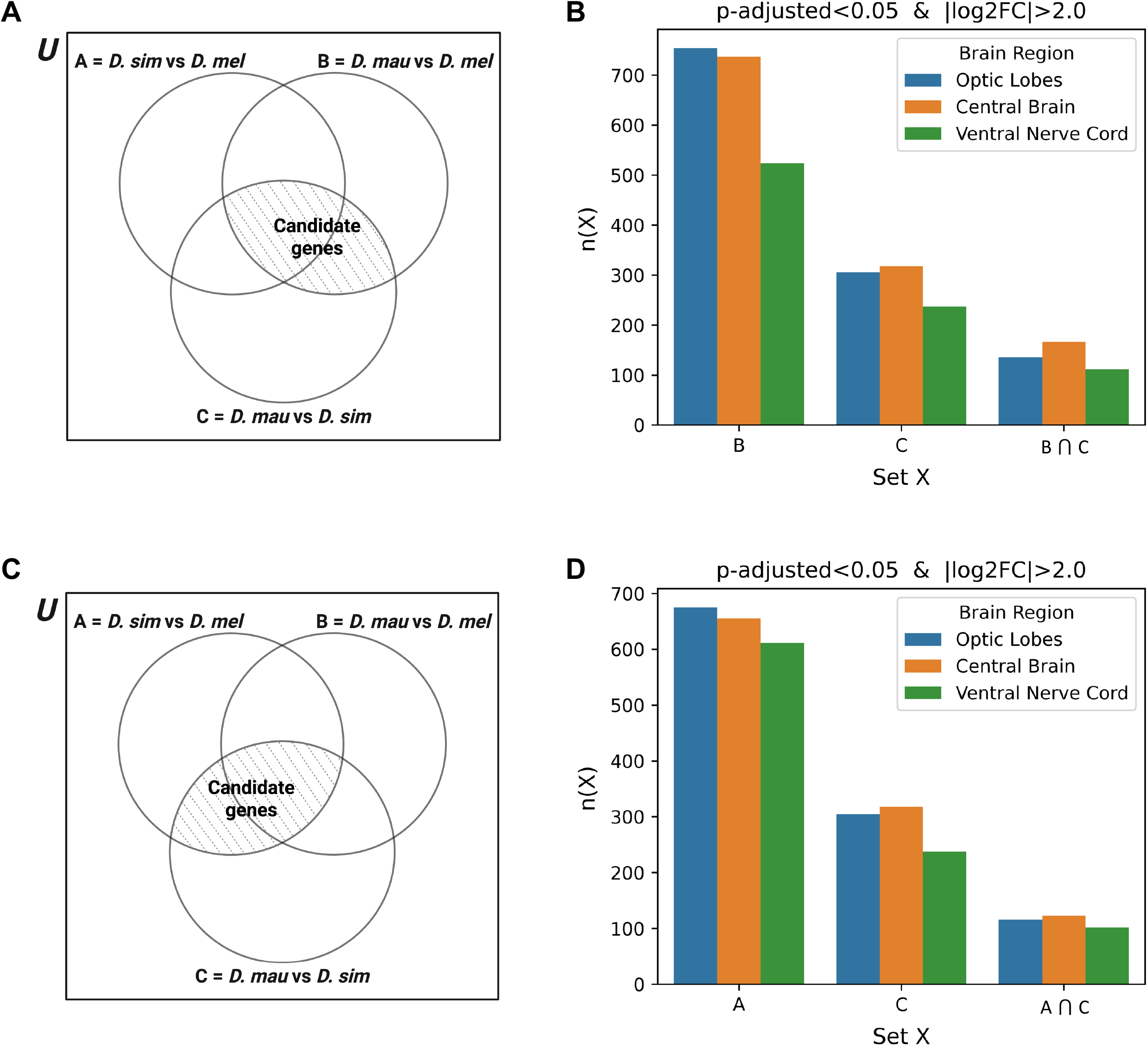
Candidate neuronal gene set sizes corresponding to different sibling species behavioral combinations. Related to Figure 3. (A)Set operations strategy for 2) in Figure 3A to constrain the number of candidate neuronal genes underlying observed inter-specific differences in locomotor activity behavior. The universal set (*U*) is defined as the finite set of all annotated genes in the *D. melanogaster* reference genome. (B)Comparison of brain region-specific candidate neuronal gene set sizes before and after executing set operations strategy in (A). n(X) denotes the cardinality of a given set X (i.e. the number of differentially expressed genes in set X). (C)Set operations strategy for 3) in Figure 3A to constrain the number of candidate neuronal genes underlying observed inter-specific differences in locomotor activity behavior. The universal set (*U*) is defined as the finite set of all annotated genes in the *D. melanogaster* reference genome. (D)Comparison of brain region-specific candidate neuronal gene set sizes before and after executing set operations strategy in (C). n(X) denotes the cardinality of a given set X (i.e. the number of differentially expressed genes in set X).

## SUPPLEMENTAL DATASET LEGENDS

**Dataset S1**. Raw read counts for all 27 bulk RNA-seq samples. Related to Figures 1C, S1, and S2.

**Dataset S2**. Number of differentially expressed genes for each pairwise species comparison per brain region, p-adjusted<0.05. Related to Figure 1D.

**Dataset S3**. Number of differentially expressed genes for each pairwise species comparison per brain region, p-adjusted<0.01. Related to Figure 1D.

**Dataset S4**. Beam transition counts for locomotor activity behavior assays. Related to Figure 2A.

**Dataset S5**. Differentially expressed genes in Optic Lobes for *D. simulans* vs *D. melanogaster*, p-adjusted<0.05 and |log_2_FC|>2.0. Related to Figure 2D.

**Dataset S6**. Differentially expressed genes in Optic Lobes for *D. mauritiana* vs *D. melanogaster*, p-adjusted<0.05 and |log_2_FC|>2.0. Related to Figure 2D.

**Dataset S7**. Differentially expressed genes in Optic Lobes for *D. mauritiana* vs *D. simulans*, p-adjusted<0.05 and |log_2_FC|>2.0. Related to Figure 2D.

**Dataset S8**. Differentially expressed genes in Central Brain for *D. simulans* vs *D. melanogaster*, p-adjusted<0.05 and |log_2_FC|>2.0. Related to Figure 2D.

**Dataset S9**. Differentially expressed genes in Central Brain for *D. mauritiana* vs *D. melanogaster*, p-adjusted<0.05 and |log_2_FC|>2.0. Related to Figure 2D.

**Dataset S10**. Differentially expressed genes in Central Brain for *D. mauritiana* vs *D. simulans*, p-adjusted<0.05 and |log_2_FC|>2.0. Related to Figure 2D.

**Dataset S11**. Differentially expressed genes in Ventral Nerve Cord for *D. simulans* vs *D. melanogaster*, p-adjusted<0.05 and |log_2_FC|>2.0. Related to Figure 2D.

**Dataset S12**. Differentially expressed genes in Ventral Nerve Cord for *D. mauritiana* vs *D. melanogaster*, p-adjusted<0.05 and |log_2_FC|>2.0. Related to Figure 2D.

**Dataset S13**. Differentially expressed genes in Ventral Nerve Cord for *D. mauritiana* vs *D. simulans*, p-adjusted<0.05 and |log_2_FC|>2.0. Related to Figure 2D.

**Dataset S14**. Concatenated lists of differentially expressed gene names from Datasets S5 to S13. Related to Figure 2D.

**Dataset S15**. Number of candidate neuronal genes per brain region for each set and after performing set operations strategy on Dataset S14. Related to Figures 2D, S3B, and S3D.

**Dataset S16**. STAR RNA-seq aligner performance statistics for all bulk RNA-seq samples. Related to Figure S2.

## SUPPLEMENTAL SOFTWARE LEGENDS

**Software S1**. R script for DESeq2 analysis and visualization of all bulk RNA-seq samples. Related to Figures 1C, S1A, and S1B.

**Software S2**. R script for DESeq2 analysis of region-specific bulk RNA-seq samples. Related to Figure 1D.

**Software S3**. Python script for plotting behavior data and statistical analysis. Related to Figure 2A.

**Software S4**. R script for map visualization of *Drosophila* species line geographical origins. Related to Figure 2B.

**Software S5**. Python script for implementing set operations strategies. Related to Figures 2D, 3A, S3B, and S3D.

