## Supplementary material for "Molecular Basis of Behavioral Diversity in a Sibling Species Trio": Software S2

### R version 4.4.2. "Molecular Basis of Behavioral Diversity in a Sibling Species Trio" (2026). Software S2. 
### This script performs differential gene expression analysis for specific brain regions using the DESeq2 package in R.
### Note that the Dataset S1 raw read counts input file contains all 27 bulk RNA-seq samples. Two pre-identified outlier 
### samples are removed by this script before running the DESeq2 analysis. 

## Load libraries
library(DESeq2)
library(janitor)
library(textshape)

## Load raw read counts file (Dataset S1)
counts <- read.delim("Dataset S1.txt", header=T, sep="\t")

## Remove two sample outlier columns
counts <- dplyr::select(counts, -c("mau_VNC_b", "sim_OL_a"))

## Remove OL and CB columns to obtain dataframe with only VNC samples (adjust based on region of interest for analysis)
counts <- dplyr::select(counts, -c("sim_CB_a", "mel_OL_a", "mel_CB_a", "mau_OL_a", "mau_CB_a",
                                  "sim_OL_b", "sim_CB_b", "mel_OL_b", "mel_CB_b", "mau_OL_b", "mau_CB_b",
                                  "sim_OL_c", "sim_CB_c", "mel_OL_c", "mel_CB_c", "mau_OL_c", "mau_CB_c"))

## Format data structure
rownames(counts) <-counts$ID
counts <- counts[,-1]
counts <- column_to_rownames(counts, loc = 1)
counts <- ceiling(counts)
counts.transposed <- t(counts)

library_info <- data.frame(ID=rownames(counts.transposed))
all(colnames(counts)==library_info$ID)
library_info$sample <- c('a', 'a', 'a', 'b', 'b', 'c', 'c', 'c')  #adjust according to brain region
library_info$species <- c('sim', 'mel', 'mau', 'sim', 'mel', 'sim', 'mel', 'mau')  #adjust according to brain region

library_info$species <- as.factor(library_info$species)
library_info$species <- factor(library_info$species, levels=c('sim', 'mel', 'mau'))


## Run DESeq2 analysis
dds <- DESeqDataSetFromMatrix(countData=counts, colData=library_info, design=~species)
dds <- DESeq(dds)


## Obtaining differentially-expressed genes that meet specified significance thresholds
res_species <- results(dds, contrast=c('species', 'mau', 'sim'))  #adjust species for each pairwise test
summary(res_species, alpha=0.05) #print number of DEGs for p-adjusted<0.05
summary(res_species, alpha=0.01) #print number DEGs for p-adjusted<0.01

res_no_na <- res_species[!is.na(res_species$log2FoldChange), ]   #remove rows where log2FoldChange value is NA
res_species_FC2 <- res_no_na[abs(res_no_na$log2FoldChange) > 2,]  #subset rows where absolute log2FoldChange value larger than 2
summary(res_species_FC2, alpha=0.05) #print number of DEGs for p-adjusted<0.05 and |log2FC|>2 
write.table(subset(res_species_FC2, padj < 0.05), file="mau_sim_VNC.deseq2.significant.txt", sep="\t") #write results to text file
