## Supplementary material for "Molecular Basis of Behavioral Diversity in a Sibling Species Trio": Software S5

### Python version 3.11.5. "Molecular Basis of Behavioral Diversity in a Sibling Species Trio" (2026). Software S5.
### This script performs set operations strategies to obtain the sets of candidate genes and their sizes for
### Drosophila sibling species trio behavior phenotype combinations 1), 2), and 3) in Figure 3A. 
### Only the VNC brain region is examined in this example, but can be substituted with CB and OL. 
### Input file is Dataset S14.


import pandas as pd

## Load dataframe of differentially expressed gene names (Dataset S14)
df = pd.read_csv('Dataset S14.csv')


## Convert columns of interest to lists
sim_mel_VNC = df['sim_mel_VNC'].values.tolist() #VNC is the region of interest here and can be substituted with OL or CB
mau_mel_VNC = df['mau_mel_VNC'].values.tolist()
mau_sim_VNC = df['mau_sim_VNC'].values.tolist()


## Convert lists to pandas Series and drop NaNs
sim_mel_VNC1 = pd.Series(sim_mel_VNC).dropna()
mau_mel_VNC1 = pd.Series(mau_mel_VNC).dropna()
mau_sim_VNC1 = pd.Series(mau_sim_VNC).dropna()


## Convert the cleaned Series back to lists
sim_mel_VNC2 = sim_mel_VNC1.tolist()
mau_mel_VNC2 = mau_mel_VNC1.tolist()
mau_sim_VNC2 = mau_sim_VNC1.tolist()

## Finding the lengths and elements of candidate gene sets using set operations strategies
# Behavior combination 1)
common_elements = set(sim_mel_VNC2) & set(mau_mel_VNC2)
cardinality_common_elements = len(common_elements)
print(f"The number of unique common genes in A and B is: {cardinality_common_elements}")

candidates = list(set(common_elements) - set(mau_sim_VNC2))
candidate_count = len(candidates)
print(f"The number of candidate genes for (AintB)\C is: {candidate_count}")


# Behavior combination 2)
common_elements = set(mau_mel_VNC2) & set(mau_sim_VNC2)
cardinality_common_elements = len(common_elements)
print(f"The number of unique common genes/candidate genes in B and C is: {cardinality_common_elements}")


# Behavior combination 3)
common_elements = set(sim_mel_VNC2) & set(mau_sim_VNC2)
cardinality_common_elements = len(common_elements)
print(f"The number of unique common genes/candidate genes in A and C is: {cardinality_common_elements}")
